# Mitochondrial DNA Sequence Diversity in Mammals: a Correlation Between the Effective and Census Population Sizes

**DOI:** 10.1101/2020.02.28.969592

**Authors:** Jennifer James, Adam Eyre-Walker

## Abstract

What determines the level of genetic diversity of a species remains one of the enduring problems of population genetics. Since, neutral diversity depends upon the product of the effective population size and mutation rate there is an expectation that diversity should be correlated to measures of census population size. This correlation is often observed for nuclear but not for mitochondrial DNA. Here we revisit the question of whether mitochondrial DNA sequence diversity is correlated to census population size by compiling the largest dataset to date from 639 mammalian species. In a multiple regression we find that nucleotide diversity is significantly correlated to both range size and mass-specific metabolic rate, but not a variety of other factors. We also find that a measure of the effective population size, the ratio of non-synonymous to synonymous diversity, is also significantly negatively correlated to both range and mass-specific metabolic rate. These results together suggest that species with larger ranges have larger effective population sizes. The slope of the relationship between diversity and range is such that doubling the range increases diversity by 12 to 20%, providing one of the first quantifications of the relationship between effective and census population sizes.

## Introduction

One of the central aims of population genetics is to understand why genetic diversity varies between species. However, despite five decades of research and the fact that nucleotide diversities vary by over two orders of magnitude (Lynch and Conery 2003; Leffler, et al. 2012), we still have a poor understanding of the factors that affect genetic diversity at the DNA level (Lewontin 1974; Leffler, et al. 2012). Since the level of neutral diversity is expected to depend upon the product of the mutation rate per generation and the effective population size, there has been an expectation that diversity should depend on the census population size. This expectation has generally been met in analyses of nuclear DNA diversity between species (see Table S1); for example, some of the earliest analyses of allozyme diversity found heterozygosity to be positively correlated to population size across species (Soule 1976; Nei and Graur 1984) and this pattern has been found in almost all subsequent analyses (Table S1). One notable exception is a recent analysis of nuclear nucleotide diversity across diverse animal species (Romiguier, et al. 2014). This is surprising, because it is by far the largest nuclear dataset collected yet in terms of loci sampled. However, (Romiguier, et al. 2014) had no direct estimate of the population size for most of their species, and for most of them it is extremely difficult to estimate; instead they used a crude estimate of population size, the distance between the two furthest sampled individuals.

Although, there is generally a positive correlation between diversity and measures of population size for nuclear DNA, diversity increases slowly relative to census population size, a pattern that has become known as Lewontin’s paradox; for example, in the early analyses of Soule (1976) and Nei and Graur (1984) they found that allozyme heterozygosity was linearly related to the logarithm of population size. However, more recent studies have not investigated the relationship between diversity and census population size quantitatively, instead just reporting whether there is a significant relationship between an estimate, or likely correlate of the census population size, and genetic diversity.

In contrast to nuclear DNA, many studies have failed to find a correlation between diversity in mitochondrial DNA and measures of population size between species (Table S2). Even when a correlation exists for the same species for nuclear DNA, a correlation for mtDNA is not necessarily observed (e.g. see Bazin et al.(2006), Singhal et al. (2017)). Bazin et al. (2006) have ascribed the lack of a correlation between mitochondrial diversity and census population size to genetic hitch-hiking, which might potentially have two effects. First, as Maynard Smith and Haigh (1974) first suggested, genetic hitch-hiking might increase in frequency as population size increases if the rate of adaptive evolution is mutation limited. As Gillespie (2000) has shown, this can lead to a disconnect between levels of diversity and population size. Second, hitch-hiking might simply increase the variance in levels of diversity, making it more difficult to observe a correlation, even if one exists.

Although, many previous analyses have generally failed to observe a relationship between diversity and measures of population size for mtDNA they have tended to either look over very broad phylogenetic scales or have limited sample size. Considering organisms over very broad phylogenetic scales might make it difficult to detect any correlation between diversity and population size because many other factors might also vary. Here we reconsider the relationship between mtDNA diversity and population size in mammals using the most extensive dataset compiled to date, and we attempt to quantify the relationship between the census and effective population sizes for the first time in a large dataset. We investigate whether diversity is correlated to a measure of census population size, the species range, but we also investigate whether it is correlated to a number of life history and demographic variables, as potential correlates of population density and the mutation rate, two other factors that might be expected to affect levels of neutral diversity.

## Results

We collected mitochondrial polymorphism data from 639 mammalian species for which at least four individuals have been sequenced. The average number of individuals sequenced was 15 and the average length of our alignments was 1300 bp. We also compiled life history and demographic information for many of these species. Variables included in the analysis were: range size, absolute latitude, adult body mass, age of sexual maturity, longevity and mass-specific metabolic rate (MSMR). These were chosen because they have either been shown to be correlated to diversity in previous analyses or might act as proxies for population density or the mutation rate. All of our variables show a significant phylogenetic signal, with Pagel’s λ close to one for everything except our two diversity statistics and range (Table 1). As a consequence, we used the method of independent contrasts in all analyses (Felsenstein 1985).

**Table 1.** Testing for phylogenetic inertia using Pagel’s λ, using log transformed data. The p-value is from a likelihood ratio test against the hypothesis that there is no phylogenetic signal, i.e. *λ* = 0.

| **Trait (log values)** | ***n*** | **Pagel's $\lambda$** | ***P*-value** |
| --- | --- | --- | --- |
| $\pi_S$ | | 0.39 | $1 \times 10^{-14}$ |
| $\pi_N / \pi_S$ | | 0.40 | $4 \times 10^{-12}$ |
| **Mass** | | 1.0 | $2 \times 10^{-306}$ |
| **Longevity** | | 0.92 | $3 \times 10^{-70}$ |
| **Sexual maturity** | | 0.95 | $2 \times 10^{-85}$ |
| **Mass-specific metabolic rate** | | 0.99 | $1 \times 10^{-31}$ |
| **Range** | | 0.64 | $4 \times 10^{-35}$ |
| **Absolute latitude** | | 0.84 | $6 \times 10^{-72}$ |

### Genetic diversity

There is little evidence that selection acts upon synonymous sites in mammalian mitochondrial DNA, and hence we use synonymous diversity as a measure of neutral genetic diversity. We find that synonymous nucleotide diversity, *π*_*S*_, is significantly positively correlated to the geographic range of a species and MSMR (Figure 1, Table 2), and negatively correlated to the absolute latitude, and age at sexual maturity (Table 2). However, many of the life history and demographic traits in mammals are correlated; we therefore used a multiple linear regression modelling approach to consider the joint effects of traits on *π*_*S*_. In the following models we exclude longevity: this variable is strongly correlated to age at sexual maturity (Pearson’s R on log transformed data = 0.82, *p* = 2×10^−47^), and unlike age at sexual maturity it is not correlated to *π*_*S*_ in a single linear regression. We are also unable to include both mass and MSMR in a single model, because they are very strongly negatively correlated (R = −0.88, *p* = 5×10^−49^), leading to high variance inflation factors. However, we decided to include MSMR as it is correlated to *π*_*S*_.

**Table 2.** Results of correlation analyses for two molecular evolutionary traits in mitochondrial DNA: *π*_*S*_ and *π*_*N*_/*π*_*S*_, with life history and demographic traits. Values are log-transformed before phylogenetic contrasts are calculated. The column *n* gives the number of contrasts available for each correlation. Significant results are in bold.

| Trait (Log values) |  | n | Pearson's correlation coefficient |  |
| --- | --- | --- | --- | --- |
|  |  |  | R | p |
| Mass | $\pi_S$ | 537 | -0.039 | 0.37 |
| | $\pi_N/\pi_S$ | 466 | 0.034 | 0.47 |
| Longevity | $\pi_S$ | 225 | -0.056 | 0.40 |
| | $\pi_N/\pi_S$ | 202 | 0.071 | 0.31 |
| Age of sexual Maturity | $\pi_S$ | 238 | -0.16 | **0.011** |
| | $\pi_N/\pi_S$ | 217 | 0.029 | 0.67 |
| Mass-specific metabolic rate | $\pi_S$ | 144 | 0.20 | **0.018** |
| | $\pi_N/\pi_S$ | 129 | -0.22 | **0.012** |
| Range | $\pi_S$ | 556 | 0.32 | **$2 \times 10^{-14}$** |
| | $\pi_N/\pi_S$ | 476 | -0.21 | **$2 \times 10^{-6}$** |
| Absolute latitude | $\pi_S$ | 556 | -0.11 | **0.013** |
| | $\pi_N/\pi_S$ | 476 | 0.060 | 0.19 |

**Figure 1.**
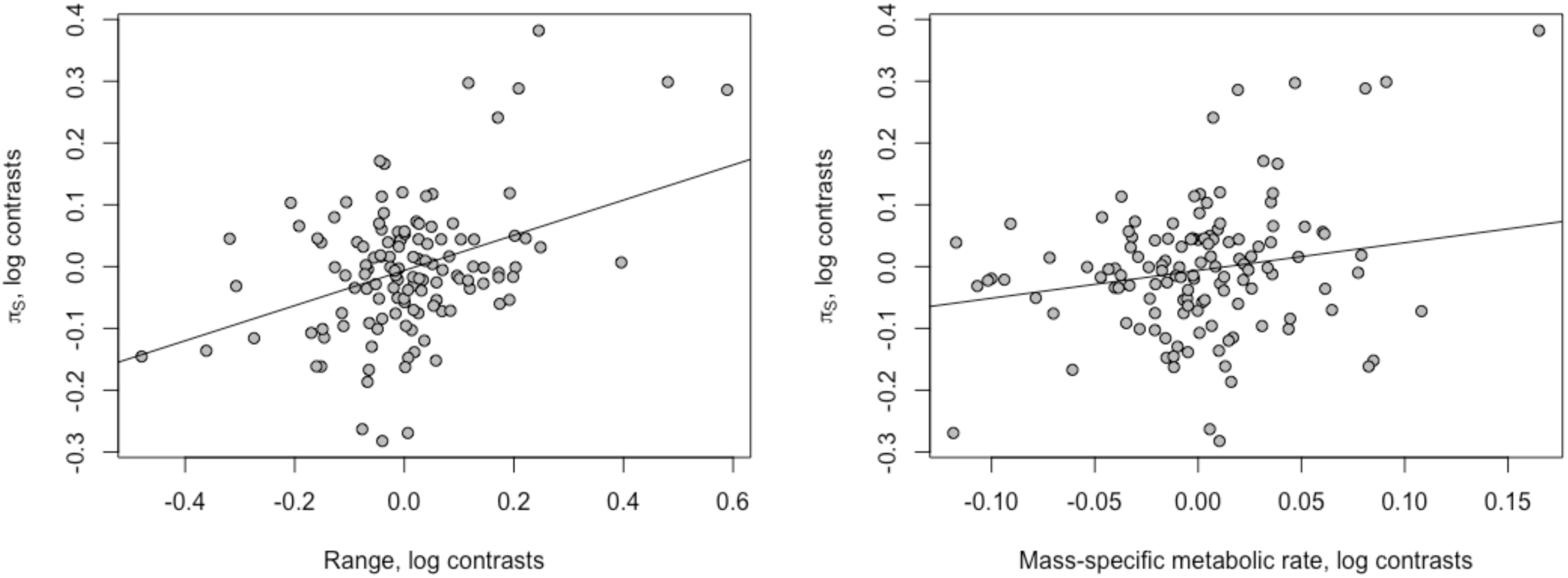
The correlation between *π*_*S*_ and its two strongest predictors: the global range of a species, and the mass-specific metabolic rate of a species. The values plotted are log-transformed phylogenetic contrasts. (The lines shown have the slope from the multiple linear regression model of *π*_*S*_, including range and mass, range slope = 0.28, *p* = 1×10^−5^, MSMR slope = 0.44, *p* = 0.024.)

We find that in a multiple linear regression of all remaining traits, only geographic species range and MSMR are significantly correlated to *π*_*S*_. Although age at sexual maturity is significantly correlated to *π*_*S*_ (see Table 1), including this variable does not significantly improve the model fit (ANOVA test on linear models with range and MSMR, and either with or without age at sexual maturity: *p* = 0.74) and if we remove this variable, our dataset increases to 128 species contrasts from 87 species contrasts. Including latitude does not significantly improve the fit of the model (*p* = 0.75). Overall, a multiple linear regression for *π*_*S*_ including range and MSMR has an overall adjusted R-squared = 0.20, *p* = 5×10^−7^, with both variables being positively correlated to *π*_*S*_.

Range size and MSMR explain relatively little of the variance in synonymous diversity – respectively 10% and 4.0% (Figure 1, Table 2). The slopes are also shallow. For range the slope between log contrast in synonymous diversity and log contrast in range is 0.16 in a simple regression and 0.28 in a multiple regression and this implies that as range size doubles, so diversity increases by just 12 or 20% respectively. For MSMR the slope is 0.54 or 0.44 suggesting that as MSMR doubles so *π*_*S*_ increases by 45% or 36%.

### The efficiency of selection

Neutral genetic diversity is expected to be a product of *N*_*e*_ and the mutation rate. It seems likely that range is a correlate of census population size and this affects the effective population size. The origins of the correlation between diversity and MSMR are less clear; it might be that species with high MSMR have high mutation rates, but it is also possible that MSMR is related to *N*_*e*_ in some way, possibly through population density. To investigate the correlation between diversity and MSMR in more depth, and to confirm that the correlation with range is driven by variation in *N*_*e*_, we investigated whether an alternative measure of *N*_*e*_, the ratio of non-synonymous to synonymous nucleotide diversity, was correlated to range, MSMR and the other variables we have considered. We find that *π*_*N*_/*π*_*S*_ is only correlated to two variables, range (r = −0.21, slope = −0.092, *p* = 2×10^−6^) and MSMR (r = −0.22, slope = −0.050, *p* = 0.012) (Table 2)(Figure 2). Note that both correlations are negative, consistent with the patterns seen for *π*_*S*_ alone; they suggest that *N*_*e*_ increases with both range size and MSMR, and this leads to an increase in *π*_*S*_ and a decrease in *π*_*N*_/*π*_*S*_. As for *π*_*S*_, we performed multiple linear regression models for *π*_*N*_/*π*_*S*_, including all life history traits; however, range and MSMR remain the only two traits that are significant explain, with an overall adjusted R-squared of 0.12, *p* = 0.00024.

**Figure 2.**
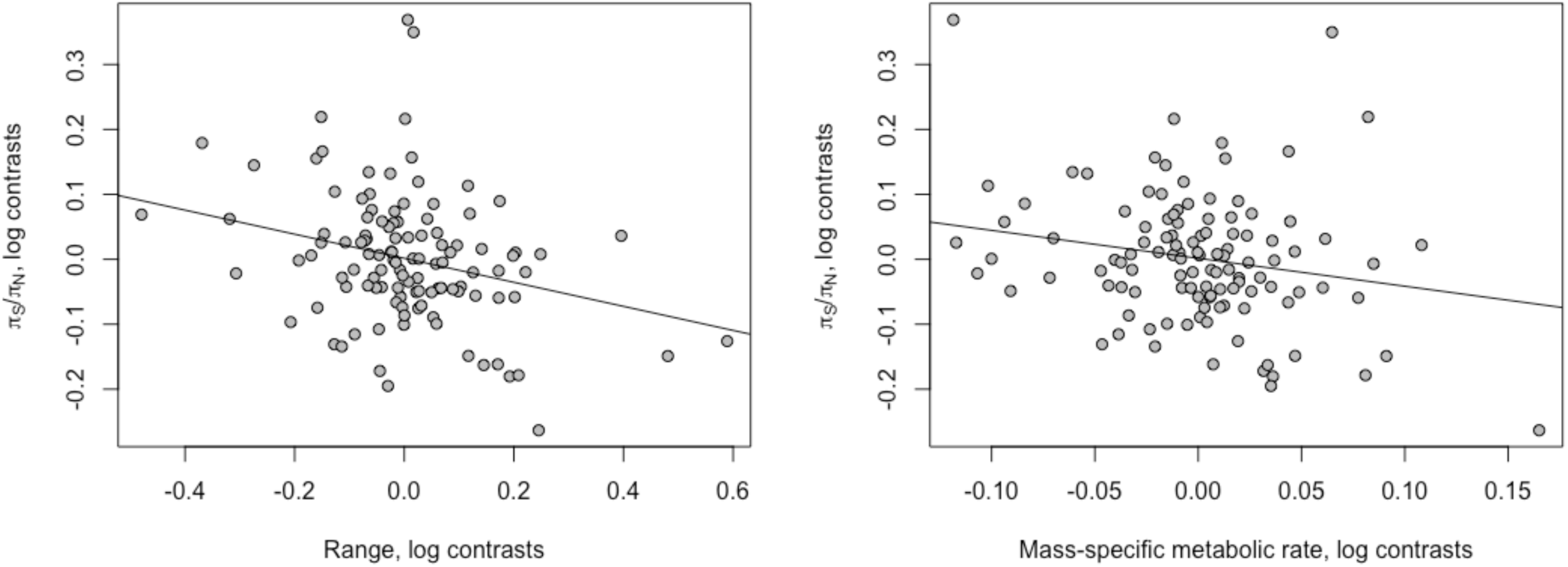
The correlation between *π*_*N*_/*π*_*S*_ and its two strongest predictors: the global range area of a species, and the mass-specific metabolic rate of a species. The values plotted are log-transformed phylogenetic contrasts. (The lines shown have the slope from the multiple linear regression model of *π*_*N*_*/π*_*S*_, including range and mass, range slope = −0.19, *p* = 0.0033, MSMR slope = −0.43, *p* = 0.033.

### Role of mutation rate

Overall, our results suggest that *N*_*e*_ is an important factor in shaping patterns of mitochondrial molecular evolution. However, mutation rate variation is also likely to affect patterns of mitochondrial diversity. To explore this, we sought to investigate whether a proxy for mutation rate, the rate of neutral divergence, *d*_*S*_, is correlated to levels of neutral genetic diversity. In this analysis, we used a sister pairs method: our dataset was divided into sets of two sister species and an outgroup, which were used to calculate divergence, and we then considered the relationship between the trait contrasts of the sister species in each set thereby correcting for phylogenetic non-independence. This dataset includes a total of 126 contrasts, however, in order to remove potentially unreliable estimates of *d*s from the dataset we excluded contrasts for which 0.00005 < *d*_*s*_ < 1 for either sister species, resulting in a dataset of 100 contrasts. The results do not qualitatively change if, instead of this exclusion step, we improve divergence estimates by including only transversion mutations. We restrict our analysis in this section to only range, body mass and latitude, the three traits for which we have the largest number of observations.

We find that *d*_*s*_ is not significantly correlated to any of our life history or demographic traits (similar results were also found by Lanfear et al. (2010)), nor to any of the polymorphism traits. Additionally, if we include *d*_*s*_ along with life history and demographic traits in a multiple linear regression model for π_s_ (*n* = 64), we find that including *d*_*s*_ does not significantly improve our model fit (ANOVA test, *p* = 0.86), indicating that *d*_*s*_ explains little variance in π_s_. Excluding *d*_*s*_, our model results are somewhat similar using this paired method as in our full PIC method as detailed above: range is possibly correlated to π_s_ (coefficient = 0.11, p = 0.096) though it is not significant. However, we do find a significant effect of latitude on π_s_ (coefficient = −0.30, p = 0.048). The overall model adjusted R-squared is 0.13, *p* = 0.011. The discrepancies may occur because in this model we have lower sample sizes and therefore likely lower power.

## Discussion

We have investigated what factors are correlated to levels of synonymous diversity in mitochondrial DNA in mammals. In a multiple regression we find that diversity is only significantly correlated to mass-specific metabolic rate and range size. Both correlations could be driven by a relationship between effective population size and census population size; species with larger ranges have larger census population sizes, and those with higher mass-specific metabolic rates have higher population density. This hypothesis is supported by the correlation between a measure of the effective population size, the efficiency of natural selection, *π*_*N*_/*π*_*S*_, and both range size and MSMR. Nevertheless, it is possible the correlations between *π*_*S*,_ *π*_*N*_/*π*_*S*_ and MSMR are a consequence of potentially two other factors. It might that organisms with higher MSMR have higher mutation rates, which leads to a positive correlation with *π*_*S*_, but also more stringent selection for metabolic efficiency, which leads to the negative correlation with *π*_*N*_/*π*_*S*_.

In a previous analysis of mitochondrial diversity in mammals Nabholz et al. (2008) found no correlation that survived phylogenetic correction, between diversity and all of the factors considered here, with the exception of latitude. Instead they found a marginally significant correlation with the substitution rate. However, their dataset was considerably smaller than ours – 179 species in their largest dataset, in which the sequences were quite short.

It is of interest to ask whether the relationship between *π*_*N*_/*π*_*S*_ and range is consistent with the relationship between *π*_*S*_ and range. Let us assume that synonymous mutations are neutral and non-synonymous mutations are deleterious with selection coefficients drawn from a gamma distributed distribution of fitness. Under this model *π*_*S*_ = 2*N*_*e*_*u* and 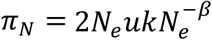, where *N*_*e*_ is the effective population size of females, *u* is the mutation rate per generation, *k* is a constant and *β* is the shape parameter of the gamma distribution (Welch et al. (Welch, et al. 2008)). Under this model, we expect the ratio of *π*_*N*_/*π*_*S*_ in species 1 relative to species 2 to equal

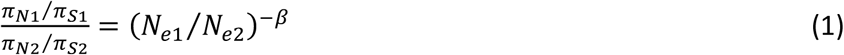

If we assume that the mutation rate and *N*_*e*_ are uncorrelated, and *N*_*e*_ is correlated to some factor x, for example range, then we can estimate the ratio of the effective population sizes from the ratio of the diversities – i.e.

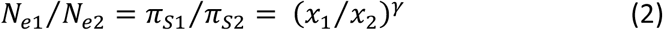

where γ is the slope of the relationship between log(*N*_*e*_) and log(range). Substituting equation 2 into equation1 gives

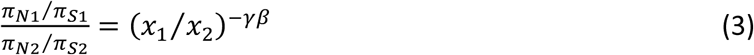

An estimate of γ can be obtained from the slope of the regression of log contrast in *π*_*S*_ against log contrast in range; for range this is 0.16. Previously we have estimated the shape parameter of the DFE to be 0.45 in mammalian mitochondrial DNA from the site frequency spectrum (James et al. 2016), and hence we predict the slope between *π*_*N*_/*π*_*S*_ and range to be −0.16 × 0.45 = −0.072, similar but slightly lower than the observed slope of −0.092. The excepted slope is less steep than the observed slope which is consistent with previous observations in both mitochondrial (James, et al. 2017) and nuclear (Chen, et al. 2017; Castellano, et al. 2018) datasets. Castellano et al. (2018) show that this is to be expected if there is genetic hitch-hiking, because hitch-hiking leads to a non-equilibrium situation in which deleterious non-synonymous genetic diversity recovers more rapidly than synonymous neutral diversity.

An important question is whether the slope of the relationship between synonymous diversity and range size reflects the true relationship between effective population size and census population size. There are several potential issues. First, if the effective population size and the mutation rate are correlated then the relationship between diversity and range will either underestimate the slope of the relationship between effective and census population sizes if *N*_*e*_ and *u* are negatively correlated, or overestimate it if they are positively correlated. We currently have little data on whether *N*_*e*_ and *u* are correlated across species for mitochondrial DNA (see limited data in Piganeau et al. (2009)), although this negative correlation is observed for nuclear DNA (Lynch 2010; Lynch, et al. 2016).

Second, range is likely to be a correlate of census population size, but population density is also important. However, so long as range and density are not themselves correlated then range should be an unbiased measure of census population size. In an attempt to get a better estimate of population size we divided the range by the product of the MSMR and body mass. Our reasoning was as follows; each unit of range only has a certain amount of energy available, so dividing the range by the basal metabolic rate of an organism should give an estimate of the population size. We find that both πS and πN/πS are more strongly correlated to this estimate of population size (πS: r = 0.43, p = 4.0 × 10^−7^, πN/πS: r = −0.35, p = 0.00016) than either is to range (Table 2). The slopes of the relationship between πS and πN/πS and the composite measure of population size are 0.28 and −0.20, steeper than we observe for range alone.

Third, the slope of the relationship between our diversity estimates and census population size may have been underestimated for statistical reasons; error in the independent variable leads to an underestimate of the slope, because very large or small values are partly due to measurement error.

Several previous analyses of nuclear DNA have also observed that whilst diversity is correlated to some measure of census population size, diversity increases less rapidly than one would expect if the effective population size was a simple function of census population size; for example, in some of the very first analyses of allozyme diversity it was found that it increased linearly as a function of log census size (Soule 1976; Nei and Graur 1984). There has been considerable debate as to why this is the case. It has been suggested that this might be a consequence of genetic hitch-hiking; as population size increases, so the rate of adaptive evolution increases, increasing the influence of hitch-hiking and hence keeping diversity in check (Maynard Smith and Haigh 1974; Gillespie 2000). For nuclear DNA this does not seem to be the case. In an analysis of data from 40 animal species Corbett-Detig et al. (2015) found evidence that hitch-hiking did tend to depress diversity more greatly in species with larger census population size, but the effect was modest; at most they estimated hitch-hiking reduced diversity by 73%. However, in mitochondrial DNA where there is little or no recombination, the effects of hitch-hiking might be more dramatic, and there is some evidence that mitochondrial DNA does undergo substantial levels of adaptive evolution in animals at least (James, et al. 2016). The second possibility is that the mutation rate is negatively correlated to the effective population size, possibly because mutation rates are driven down to a limit set by the power of genetic drift (Lynch 2010; Lynch, et al. 2016). However, this hypothesis predicts that diversity should be consistent across species, at least of similar complexity, and this is not observed. We find no evidence that a measure of the mutation rate, the synonymous divergence, is correlated to a measure of the effective population size. We are therefore not much closer to understanding why diversity scales as it does with the census population size, although in this analysis we present one of the best estimates of the quantitative relationship between effective and census population sizes.

## Materials and Methods

Mitochondrial coding DNA sequences were downloaded from Mampol, a database of mammalian polymorphisms (Egea, et al. 2007). Only those species for which there was a minimum of four sequenced individuals were included in this study. Sequences were aligned by eye using Geneious version 7.0.6 (Kearse, et al. 2012). Where multiple genes were sequenced for a single species, sequences were concatenated to produce longer alignments. Alignments were then analysed using our own scripts in order to calculate synonymous nucleotide site diversity, *π*_*S*_, and a measure of the efficiency of selection, *π*_*N*_/*π*_*S*_. This ratio is undefined if a species has no synonymous diversity, and thus such species were excluded from the analyses of *π*_*N*_/*π*_*S*_.

We added life history and demographic information to the species in our dataset by using the panTHERIA database (Jones, et al. 2009). In this analysis we focused on 6 traits: adult body mass (in g); maximum longevity (in months); age at sexual maturity (in days); geographic range (in km^2^); median latitude of geographic range, which was first converted to its absolute value such that in our dataset a relationship with latitude represents a relationship with distance from the equator; and mass specific metabolic rate, which was calculated by dividing basal metabolic rate (measured in ml O^2^/hr) by the mass (in g) of the individual from which the metabolic rate measurement was taken.

Species cannot be considered as statistically independent datapoints, due to shared ancestry. In order to remove the effects of phylogenetic non-independence from our dataset, we used the method of independent contrasts (Felsenstein 1985). All life history and molecular evolution traits were log transformed, and then phylogenetic contrasts were calculated using the *ape* package in R (Paradis, et al. 2004). Our dataset using this method has *n* – 1 contrasts, where *n* is the number of species in the dataset. The phylogenetic trees used in this study were created using *TimeTree* (Kumar, et al. 2017). All analyses were conducted in R. Graphs were created using base R, and the package *jtools*. We also quantified the level of phylognetic signal in our dataset using Pagel’s λ (reviewed in (Freckleton, et al. 2002; Kamilar and Cooper 2013)), which was calculated with the R package *phylosignal* (Keck, et al. 2016).

We also included species divergence data in our results: in order to perform this analysis, we grouped species into triplets, consisting of two sister species, more closely related to each other than any other species in the dataset, and an outgroup. To be included in this dataset, sister pair species and the outgroup had to have the same mitochondrial gene sequenced, therefore this step reduced the size of the dataset considerably. The sequences for each triplet were aligned as before, and then divergence data was calculated using PAML (Yang 2007). In the subsequent analysis, we controlled for phylogenetic effects by conducting our analyses on the relative difference in values between sister species-i.e. we calculated species1_(trait)_ / species2_(trait)_ for every trait for each sister pair, and then considered the relationships between these contrasts. Therefore, the size of our dataset using this method is determined by the number of contrasts available, not the number of overall species in the dataset. However, as some of the species included in our analysis are relatively divergent, and because mitochondrial mutation rates are very high, our estimates of substitution rates may be unreliable due to the occurrence of sites which are likely to have been hit many times by mutations.

## Acknowledgements

Funding for this study was provided by the University of Sussex and NERC, grant number NE/L502042/1.

---

**Table S1.**
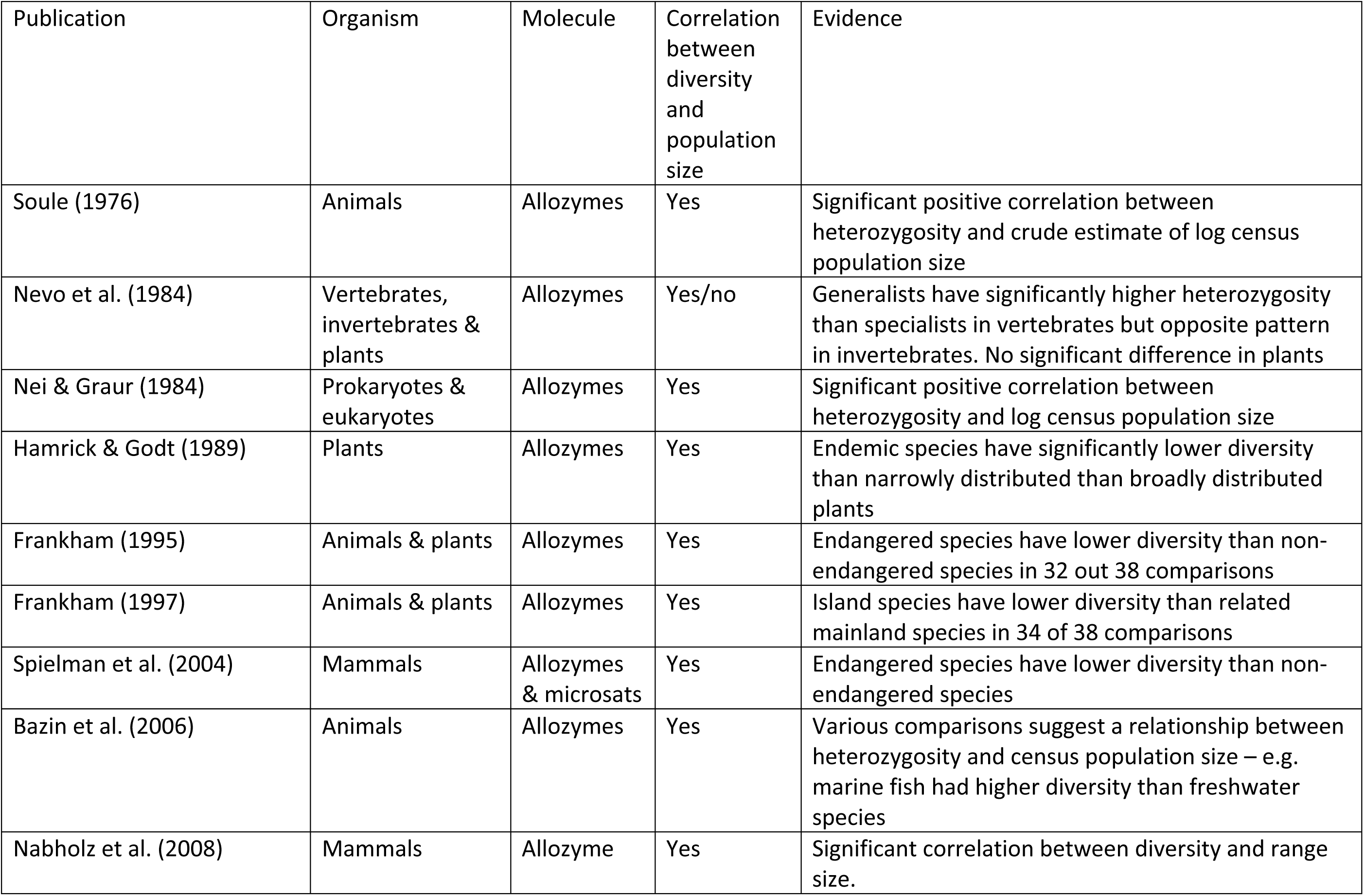
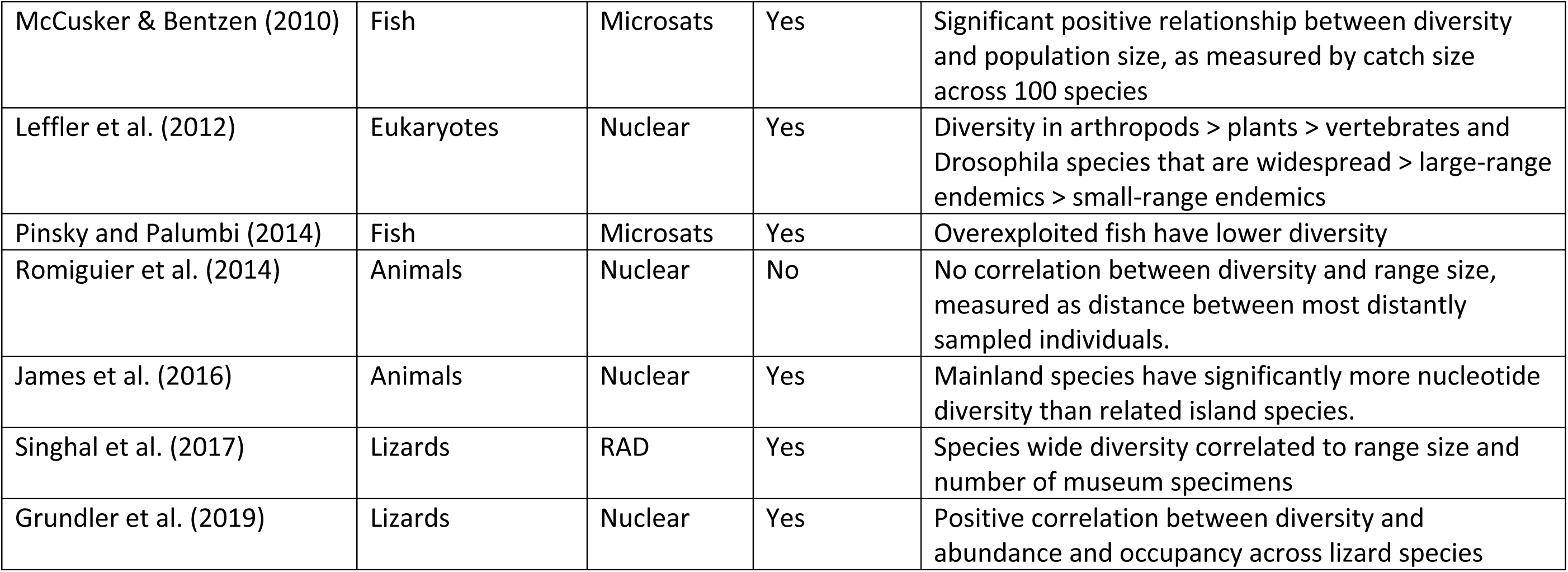
The relationship between genetic diversity and population size between species for nuclear DNA.

| Publication | Organism | Molecule | Correlation between diversity and population size | Evidence |
| --- | --- | --- | --- | --- |
| Soule (1976) | Animals | Allozymes | Yes | Significant positive correlation between heterozygosity and crude estimate of log census population size |
| Nevo et al. (1984) | Vertebrates, invertebrates & plants | Allozymes | Yes/no | Generalists have significantly higher heterozygosity than specialists in vertebrates but opposite pattern in invertebrates. No significant difference in plants |
| Nei & Graur (1984) | Prokaryotes & eukaryotes | Allozymes | Yes | Significant positive correlation between heterozygosity and log census population size |
| Hamrick & Godt (1989) | Plants | Allozymes | Yes | Endemic species have significantly lower diversity than narrowly distributed than broadly distributed plants |
| Frankham (1995) | Animals & plants | Allozymes | Yes | Endangered species have lower diversity than non-endangered species in 32 out 38 comparisons |
| Frankham (1997) | Animals & plants | Allozymes | Yes | Island species have lower diversity than related mainland species in 34 of 38 comparisons |
| Spielman et al. (2004) | Mammals | Allozymes & microsats | Yes | Endangered species have lower diversity than non-endangered species |
| Bazin et al. (2006) | Animals | Allozymes | Yes | Various comparisons suggest a relationship between heterozygosity and census population size – e.g. marine fish had higher diversity than freshwater species |
| Nabholz et al. (2008) | Mammals | Allozyme | Yes | Significant correlation between diversity and range size. |
| McCusker & Bentzen (2010) | Fish | Microsats | Yes | Significant positive relationship between diversity and population size, as measured by catch size across 100 species |
| Leffler et al. (2012) | Eukaryotes | Nuclear | Yes | Diversity in arthropods > plants > vertebrates and Drosophila species that are widespread > large-range endemics > small-range endemics |
| Pinsky and Palumbi (2014) | Fish | Microsats | Yes | Overexploited fish have lower diversity |
| Romiguier et al. (2014) | Animals | Nuclear | No | No correlation between diversity and range size, measured as distance between most distantly sampled individuals. |
| James et al. (2016) | Animals | Nuclear | Yes | Mainland species have significantly more nucleotide diversity than related island species. |
| Singhal et al. (2017) | Lizards | RAD | Yes | Species wide diversity correlated to range size and number of museum specimens |
| Grundler et al. (2019) | Lizards | Nuclear | Yes | Positive correlation between diversity and abundance and occupancy across lizard species |

**Table S2.**
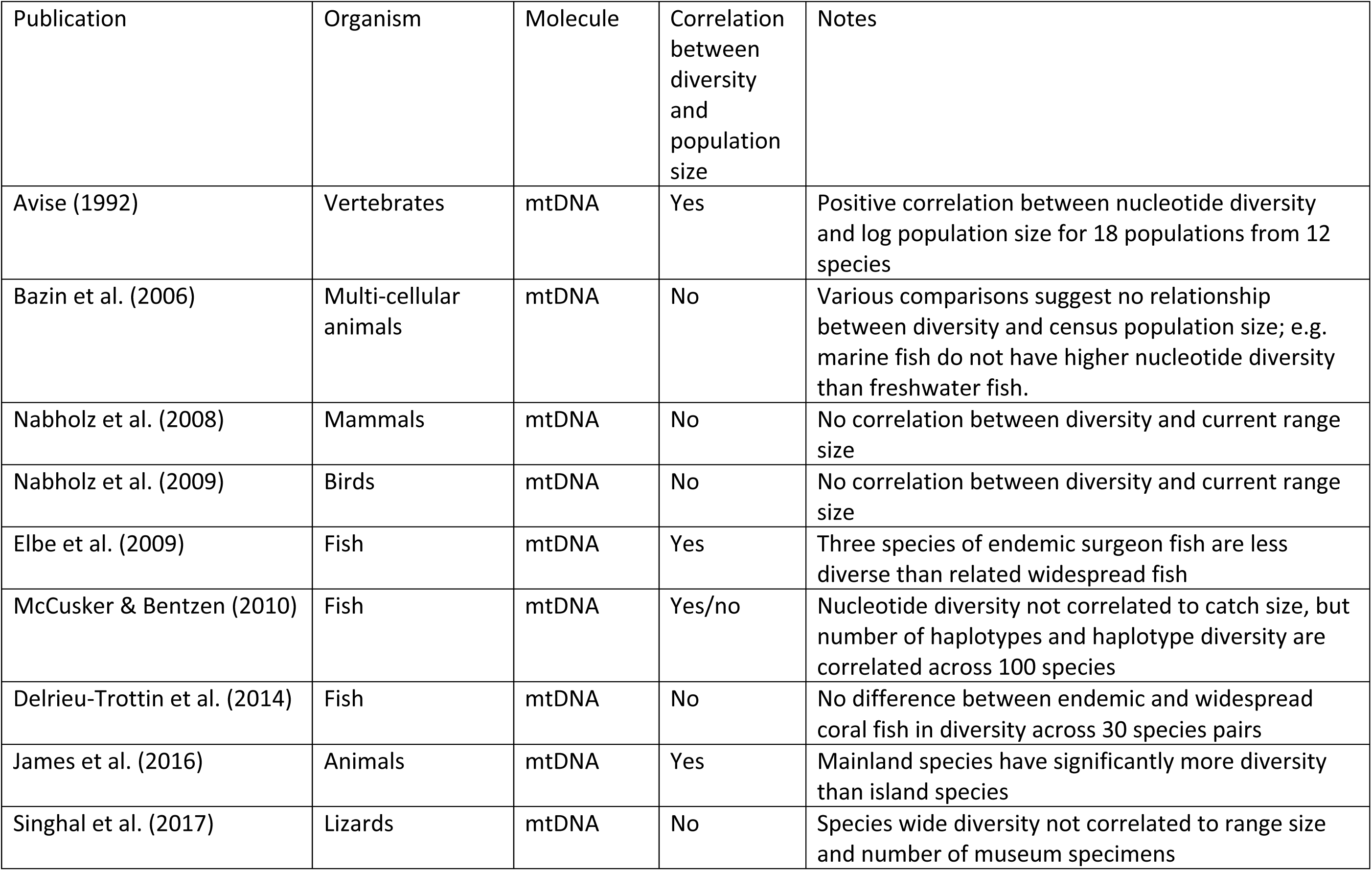
The relationship between genetic diversity and population size between species for mitochondrial DNA.

## References

Bazin E, Glemin S, Galtier N. 2006. Population size does not influence mitochondrial genetic diversity in animals. Science 312:570–572.

Castellano D, James J, Eyre-Walker A. 2018. Nearly Neutral Evolution across the Drosophila melanogaster Genome. Mol Biol Evol 35:2685–2694.

Chen J, Glemin S, Lascoux M. 2017. Genetic Diversity and the Efficacy of Purifying Selection across Plant and Animal Species. Mol Biol Evol 34:1417–1428.

Corbett-Detig RB, Hartl DL, Sackton TB. 2015. Natural selection constrains neutral diversity across a wide range of species. PLoS Biol 13:e1002112.

Egea R, Casillas S, Fernandez E, Senar MA, Barbadilla A. 2007. MamPol: a database of nucleotide polymorphism in the Mammalia class. Nucleic Acids Res 35:D624–629.

Felsenstein J. 1985. Phylogenies and the comparative method. Am. Nat. 125:1–15.

Freckleton RP, Harvey PH, Pagel M. 2002. Phylogenetic analysis and comparative data: A test and review of evidence. American Naturalist 160:712–726.

Gillespie JH. 2000. Genetic drift in an infinite population: the pseudohitchhiking model. Genetics 155:909–919.

James J, Castellano D, Eyre-Walker A. 2017. DNA sequence diversity and the efficiency of natural selection in animal mitochondrial DNA. Heredity (Edinb) 118:88–95.

James JE, Piganeau G, Eyre-Walker A. 2016. The rate of adaptive evolution in animal mitochondria. Mol Ecol 25:67–78.

Jones KE, Bielby J, Cardillo M, Fritz SA, O’’Dell J, Orme CDL, Safi K, Sechrest W, Boakes EH, Carbone C, et al. 2009. PanTHERIA: A Species-Level Database of Life History, Ecology, and Geography of Extant and Recently Extinct Mammals. Ecology 90:2648–2648.

Kamilar JM, Cooper N. 2013. Phylogenetic signal in primate behaviour, ecology and life history. Philosophical Transactions of the Royal Society B-Biological Sciences 368.

Kearse M, Moir R, Wilson A, Stones-Havas S, Cheung M, Sturrock S, Buxton S, Cooper A, Markowitz S, Duran C, et al. 2012. Geneious Basic: An integrated and extendable desktop software platform for the organization and analysis of sequence data. Bioinformatics 28:1647–1649.

Keck F, Rimet F, Bouchez A, Franc A. 2016. phylosignal: an R package to measure, test, and explore the phylogenetic signal. Ecology and Evolution 6:2774–2780.

Kumar S, Stecher G, Suleski M, Hedges SB. 2017. TimeTree: A Resource for Timelines, Timetrees, and Divergence Times. Molecular Biology and Evolution 34:1812–1819.

Lanfear R, Ho SYW. 2010. Longevity, mutation rates, and the evolution of avian mitochondrial DNA. Mitochondrial DNA 21:2–2.

Leffler EM, Bullaughey K, Matute DR, Meyer WK, Segurel L, Venkat A, Andolfatto P, Przeworski M. 2012. Revisiting an old riddle: what determines genetic diversity levels within species? PLoS biology 10:e1001388.

Lewontin RC. 1974. The genetic basis of evolutionary change. New York: Columbia University Press.

Lynch M. 2010. Evolution of the mutation rate. Trends Genet 26:345–352.

Lynch M, Ackerman MS, Gout JF, Long H, Sung W, Thomas WK, Foster PL. 2016. Genetic drift, selection and the evolution of the mutation rate. Nat Rev Genet 17:704–714.

Lynch M, Conery JS. 2003. The origins of genome complexity. Science 302:1401–1404.

Maynard Smith J, Haigh J. 1974. The hitch-hiking effect of a favourable gene. Genet. Res. 23:23–35.

Nabholz B, Mauffrey JF, Bazin E, Galtier N, Glemin S. 2008. Determination of mitochondrial genetic diversity in mammals. Genetics 178:351–361.

Nei M, Graur D. 1984. Extent of protein polymorphism and the neutral mutation theory. Evol. Biol. 17:73–118.

Paradis E, Claude J, Strimmer K. 2004. APE: Analyses of Phylogenetics and Evolution in R language. Bioinformatics 20:289–290.

Piganeau G, Eyre-Walker A. 2009. Evidence for variation in the effective population size of animal mitochondrial DNA. PLoS ONE 4:e4396.

Romiguier J, Gayral P, Ballenghien M, Bernard A, Cahais V, Chenuil A, Chiari Y, Dernat R, Duret L, Faivre N, et al. 2014. Comparative population genomics in animals uncovers the determinants of genetic diversity. Nature 515:261–U243.

Singhal S, Huang HT, Title PO, Donnellan SC, Holmes I, Rabosky DL. 2017. Genetic diversity is largely unpredictable but scales with museum occurrences in a species-rich Glade of Australian lizards. Proceedings of the Royal Society B-Biological Sciences 284.

Soule ME. 1976. Allozyme variation, its determinants in space and time. In: Ayala F, editor. Molecular Evolution. Sunderland, Massachusetts: Sinauer Associates. p. 60–77.

Welch JJ, Eyre-Walker A, Waxman D. 2008. Divergence and Polymorphism Under the Nearly Neutral Theory of Molecular Evolution. J Mol Evol 67:418–426.

## References

Avise JC. 1992. Molecular population structure and the biogeographic history of a regional fauna: a case history with lessons for conservation biology. Oikos 63:62–76.

Delrieu-Trottin E, Planes S, Williams JT. 2014. Endemic and widespread coral reef fishes have similar mitochondrial genetic diversity. Proc. Roy. Soc. Ser. B 281.

Eble JA, Toonen RJ, Bowen BW. 2009. Endemism and dispersal: comparative phylogeography of three surgeonfishes across the Hawaiian Archipelago. Marine Biology 156:689–698.

Frankham R. 1997. Do island populations have less hgenetic variation than mainland populations? Heredity 78:311–327.

Frankham R. 1995. Effective population size/adult population size ratios in wildlife: a review. Genet Res 66:95–107.

Grundler MR, Singhal S, Cowan MA, Rabosky DL. 2019. Is genomic diversity a useful proxy for census population size? Evidence from a species-rich community of desert lizards. Mol Ecol 28:1664–1674.

Hamrick JL, Godt MJW. 1989. Allozyme diversity in plant species. In: Brown AHD, Clegg MT, Kahler AL, Weir BS, editors. Plant population genetics, breeding and genetic resources. Sunderland, Massachusetts, USA: Sinauer. p. 43–63.

James JE, Lanfear R, Eyre-Walker A. 2016. Molecular Evolutionary Consequences of Island Colonization. Genome Biol Evol 8:1876–1888.

McCusker MR, Bentzen P. 2010. Positive relationships between genetic diversity and abundance in fishes. Mol Ecol 19:4852–4862.

Nabholz B, Glemin S, Galtier N. 2009. The erratic mitochondrial clock: variations of mutation rate, not population size, affect mtDNA diversity across birds and mammals. BMC Evol Biol 9:54.

Nevo E, Beiles A, Ben-Shlomo R. 1984. The evolutionary significance of genetic diversity: ecological, demographic and life histroy correlates. In: Mani GS, editor. Evolutionary Dynamics of Genetic Diversity. Berlin: Spinger-Verlag. p. 13–213.

Pinsky ML, Palumbi SR. 2014. Meta-analysis reveals lower genetic diversity in overfished populations. Mol Ecol 23:29–39.

Spielman D, Brook BW, Frankham R. 2004. Most species are not driven to extinction before genetic factors impact them. Proc. Natl. Acad. Sci. USA 101:15261–15264.

